# Defining cat mitogenome variation and accounting for numts via multiplex amplification and Nanopore sequencing

**DOI:** 10.1101/2023.06.19.545568

**Authors:** Emily Patterson, Gurdeep Matharu Lall, Rita Neumann, Barbara Ottolini, Federico Sacchini, Aiden P. Foster, Mark A. Jobling, Jon H. Wetton

## Abstract

Hair shed by domestic cats is a potentially useful source of forensic evidence. Analysable hair DNA is predominantly mitochondrial, but the recent domestication history of cats means that mtDNA diversity is low. A 402-bp control region segment is usually sequenced, defining only a small number of distinct mitotypes in populations. Previously, we used a long-amplicon approach to sequence whole mitogenomes in a sample of blood DNAs from 119 UK cats, greatly increasing observed diversity and reducing random match probabilities. To exploit this variation for forensic analysis, we here describe a multiplex system that amplifies the cat mitogenome in 60 overlapping amplicons of mean length 360 bp, followed by Nanopore sequencing. Variants detected in multiplex sequence data from hair completely mirror those from long-amplicon data from blood from the same individuals. However, applying the multiplex to matched blood DNA reveals additional sequence variants which derive from the major feline nuclear mitochondrial insertion sequence (numt), which covers 7.9 kb of the 17-kb mitogenome and exists in multiple tandem copies. We use long-amplicon Nanopore sequencing to investigate numt variation in a set of cats, together with an analysis of published genome sequences, and show that numt arrays are variable in both structure and sequence, thus providing a potential source of uncertainty when nuclear DNA predominates in a sample. Forensic application of the test was demonstrated by matching hairs from a cat with skeletal remains from its putative mother, both of which shared a globally common mitotype at the control region. The random match probability (RMP) in this case with the CR 402-bp segment was 0.21 and this decreased to 0.03 when considering the whole mitogenome. The developed multiplex and sequencing approach, when applied to cat hair where nuclear DNA is scarce, can provide a reliable and highly discriminating source of forensic genetic evidence. The confounding effect of numt co-amplification in degraded samples where mixed sequences are observed can be mitigated by variant phasing, and by comparison with numt sequence diversity data, such as those presented here.

## Introduction

Domestic cats are among the most common household pets: in the United Kingdom, for example, there are an estimated total of about 11 million, residing in 26% of homes [1]. Within such environments, cat hairs are continuously shed and transfer readily to the belongings and clothing of associated humans. The recovery of cat hairs from a crime scene may therefore provide important evidence, linking a suspect and a victim, for instance [2, 3]. As shed hairs normally originate from the cat’s undercoat, they provide minimal diagnostic phenotypic characteristics and are of limited value in microscopic comparison [4]. Comparative visual analysis is further complicated by extensive variation of hairs even within a single animal [5].

Short tandem repeat (STR) profiling multiplexes [6, 7] have been developed for the identification of individual cats, and can be used to exploit the forensic value of shed cat hairs within an investigation [8]. Work has also been undertaken to develop single nucleotide polymorphism (SNP) panels for individual identification, and for prediction of the phenotypic characteristics of a given cat [9]. However, all these analyses depend upon nuclear DNA, which is typically at very low levels in shed hairs, since these lack the root tag [5]. During hair shaft development, both nuclear and mitochondrial DNA (mtDNA) undergo degradation, but the higher copy of mtDNA and the mitochondrion’s protective double membrane mean that mtDNA is more likely to be recoverable [10]. However, due to its maternal inheritance and apparent lack of recombination this molecule is not as informative or as discriminating as markers within the nuclear genome [11].

The 17-kb feline mitochondrial genome (mitogenome; Figure 1a) encodes 13 proteins, two ribosomal RNAs, and 22 transfer RNAs, and also contains a non-coding and therefore more variable control region (CR) [12]. Traditionally, the most studied mtDNA region for individualisation purposes is a 402-bp (coordinates: 16,814–206; Figure 1a) segment within the CR [3, 5, 13-17]. This segment is flanked by two distinct repetitive sequence sites, RS2 and RS3 [5], which are reported to display high levels of heteroplasmy, and are generally avoided in analysis [18]. Focusing on the 402-bp region has also provided the practical advantage of avoiding a widespread feline nuclear mitochondrial insertion sequence (numt; Figure 1b) which has been found in domestic, wild, and sand cats [12, 19]. This numt begins within the 3’ end of the CR (Figure 1a), is 7.9 kb in length, and in domestic cats has been estimated to be tandemly repeated 38-76 times [19].

**Figure 1:**
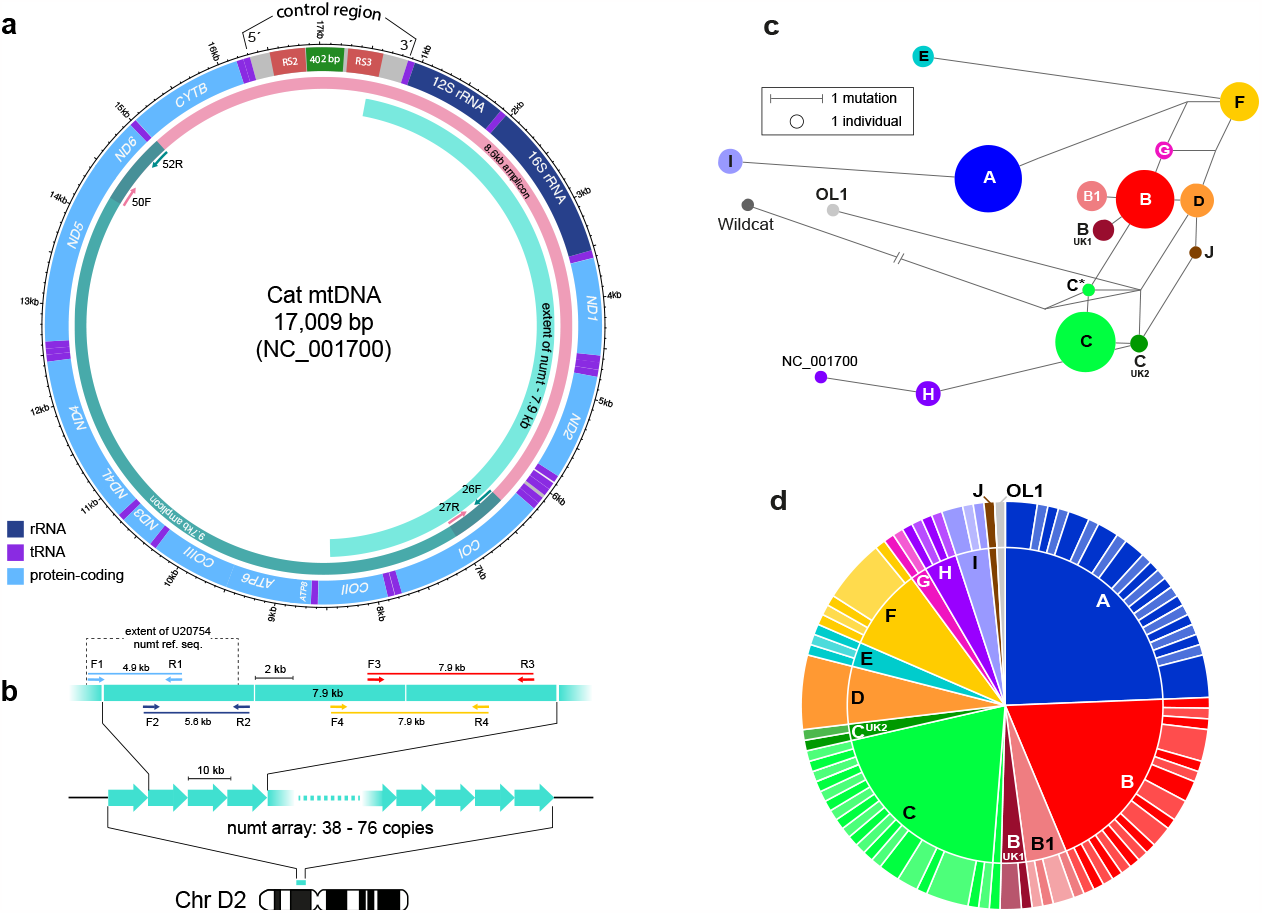
Organisation of the domestic cat mitogenome and numt array, and diversity of cat mitotypes. a) Structure of cat mtDNA based on the NC_001700 reference sequence [12], showing genes and the extent of the large numt. Primers for the two long overlapping amplicons used for sequencing [20] are indicated by short coloured arrows. The control region includes the positions of the RS2 and RS3 repeat arrays, and the 402-bp segment usually sequenced; its 5’ - 3’ orientation is according to [41]. The extent of the U20754 numt reference sequence [19] is indicated. b) Organisation and copy number range [19] of the large numt array, located on the proximal short arm of chromosome D2 [42]. Approximate positions of four primer pairs (Table S2) used to amplify segments of the numt array are indicated by short coloured arrows, with the approximate amplicon sizes also given. Note that each primer site is expected to exist within each 7.9-kb repeat unit, and (with the exception of amplicon 2) the resulting amplicon spans two adjacent repeat units in the array. c) Median-joining network based on variants within the control region within 119 cat mitogenomes [20]. Circles represent haplotypes, with area proportional to sample size, and lines between haplotypes represent mutational steps as shown in the key. The long branch to the wildcat haplotype is shortened for display. d) Pie-chart showing (inner) the diversity of 119 UK cat mitotypes as defined by the 402-bp CR sequence, and (outer) the increased diversity revealed by long-amplicon-based mitogenome sequencing (excluding the RS2 and RS3 repeat regions).

Given the uniparental mode of inheritance of mtDNA, many matrilineally related individuals are expected to share the same mtDNA haplotype (mitotype). In order to determine the evidential weight of a mitotype, its frequency should be assessed within a reference database large enough to reflect the genetic diversity present in the population [4]. Reference databases of cats sampled in different geographic locations have shown that mitotype frequencies can vary markedly [3, 5, 13, 14, 16, 17]. Based on analysis of the 402-bp CR segment, twelve ‘universal’ mitotypes designated A-L have been identified, with the four mitotypes A-D representing 60-70% of cat populations worldwide [13]. Indeed, within the UK more than half of cats shared one of two mitotypes (A or C) leading to a high observed random match probability of 0.19, based on a sample of 152 cats [3]. Figure 1c shows the frequencies and relationships of UK cat CR mitotypes [20].

To provide a higher degree of discrimination, a more powerful method than CR sequencing is necessary. We approached this by using Nanopore and Illumina technology to sequence the whole mitogenomes of 119 cats, using a long-amplicon approach to avoid the problem of the major cat numt [20]. Excluding the RS2 and RS3 repetitive regions, this allowed the extensive subdivision of mitotypes (Figure 1d), and a near order-of-magnitude reduction of random match probability of 0.16 to 0.018. Nanopore sequencing also allowed variation within the RS2 repeat array to be defined, revealing complex and dynamic minisatellite-like structures with both repeat number and repeat sequence variation, that further increase the diversity of cat mitogenomes [20].

Whole mitogenome approaches can thus offer greatly increased discrimination among domestic cats; however, long-amplicon approaches are not well suited to forensic analysis, especially in hairs, where mtDNA is expected to be degraded. The aim of this study is to develop a rapid and sensitive multiplex method to analyse cat mitogenome variation from shed hairs using short amplicons via Nanopore sequencing technology. As a case study, we apply the new method to demonstrate a mitogenome sequence match between degraded skeletal remains and hair from a matrilineally related cat.

## Materials and Methods

### Sample collection

This research was approved by the University of Leicester’s Animal Welfare and Ethical Review Body (ref.: AWERB/2021/159). Matched blood and hair samples from domestic cats were obtained by the Bristol Veterinary School. Ten domestic cat blood samples used in numt sequencing were supplied by IDEXX, as described previously [3]. Blood was dispatched in EDTA tubes at ambient temperature and frozen on arrival, and hair was placed in clip-seal plastic bags and stored at ambient temperature, between 2001 and 2021.

The disappearance of a UK-based female domestic cat in late spring 2021, and the discovery of a cat mandible eight weeks later in the same geographical area, led to the owner seeking to determine if the remains belonged to their missing cat. Both the mandible and a hair sample collected from the missing cat’s male offspring were stored at room temperature before analysis.

### DNA extraction

DNA was extracted from 200 μl of blood using the QIAamp DNA Mini Kit (Qiagen) following the manufacturer’s protocol, quantified using the NanoDrop 2000 (Thermo Scientific) and stored at -20°C.

For hair DNA extraction, 20-30 hairs per cat were washed twice in 500 μl 70% (v/v) ethanol and 1 ml Milli-Q® water. Hairs were digested with 500 μl of Buffer ATL (Qiagen), 20 μl proteinase K (20mg/mL; Qiagen) and 20 μl 0.4 M DTT (Sigma-Aldrich) at 56°C. Following overnight incubation, 40 μl proteinase K and 40 μl 0.4 M DTT were added, and the sample was incubated for a further two hours. DNA was extracted using the QIAamp DNA Mini Kit following the manufacturer’s body fluid protocol and quantified with the Qubit fluorometer 2.0 (Invitrogen) using the Qubit dsDNA HS Assay kit (Invitrogen).

In the missing cat case, a tooth was extracted from the recovered cat mandible and its crown removed. The root (∼60 mg) was incubated for 2 h at 56°C with 950 μl 0.5 M EDTA pH 8.0 (Thermo Fisher), 50 μl 1% (w/v) n-lauroylsarcosine (Sigma-Aldrich), 14 μl 1 M DTT (Sigma-Aldrich), 32 μl proteinase K (Qiagen) [21]. The sample was centrifuged at 1120 g for 3 min, and the solution then concentrated using the MinElute PCR Purification kit (Qiagen) and DNA eluted in 60 μl. The extracted DNA was quantified using the Qubit dsDNA HS Assay kit.

### Multiplex design, amplification and sequencing of mtDNA

Primer pairs were designed to cover the cat mitogenome in sixty overlapping amplicons of average length ∼360 bp. Design was based on an alignment of 119 domestic cat mitogenomes [20], with 92 complete sequences obtained through Nanopore sequencing and 26 through Illumina sequencing, plus the domestic cat reference mitogenome (NC_001700 [12]). Primers were designed with the aid of AliView (v1.26) [22] (Table S1), allowing mitotype-specific SNPs to be taken into account either by primer positioning or the incorporation of degenerate nucleotides. Potential primer interactions were assessed using AutoDimer [23].

Sixty amplicons were amplified as two sets of thirty non-overlapping amplicons each, in two separate multiplex PCRs. Two domestic cats of different haplogroups (B29 [hg A] and B30 [hg C]) were used for multiplex sequencing. Matched blood and hair samples were amplified using the Type-it Microsatellite PCR kit (Qiagen) in a 25-μl volume containing 12.5 μl of 2 x Type-It Master Mix, 0.2 μM of each primer and 0.05 ng of template DNA. Amplification conditions consisted of an initial denaturation at 95°C for 5 min, followed by 35 cycles of 95°C for 30 s, 55°C for 90 s, 72°C for 30 s, and a final extension at 68°C for 10 min. PCR products were purified using 1.8x volume of AppMag PCR Clean up Beads (Appleton Woods Ltd.) following the manufacturer’s recommendations. Equimolar concentrations of each multiplex were then pooled together for each individual’s hair or blood sample. The four samples were prepared for sequencing using the native barcoding expansion (EXP-NBD104) and ligation sequencing kit (SQK-LSK109). A 15-ng library was loaded onto a MinION flow cell (FLO-MIN106, R9.4.1) which was used for sequencing with the software MinKNOW (v20.10.3). The mitogenomes of the two individuals were also Nanopore sequenced using a long-amplicon approach (B29: ref. [20]; B30: Supplementary Data), allowing a comparison of the detected variants.

Multiplex reactions were undertaken as above with DNA extracted from the cat tooth, and from the missing cat’s offspring’s hair. Reaction products were again purified using 1.8x volume AppMag PCR Clean up Beads following the manufacturer’s instructions. Equimolar concentrations of each multiplex were pooled together for each of the two samples. The native barcoding genomic DNA protocol for Flongle (version: NBE_9065_v109_revY_14Aug2019) was followed with some minor changes. The NEBNext FFPE DNA repair mix and buffer were omitted during the DNA repair and end-prep stage where DNA libraries were prepared using 200 ng of each sample. A final library of 28 ng (∼125 fmol) was loaded onto the Flongle flow cell (FLO-FLG001) and sequencing was carried out using MinKNOW (v20.10.3) software.

### Targeted numt sequencing

NCBI’s primer-BLAST tool [24] was used to design primers specific to the major feline numt (7946 bp), which is present as a tandem array (Figure 1b; GenBank Acc: U20754; [12, 19]). The numt was amplified in four overlapping amplicons through four separate PCRs, using DNA extracted from blood samples from ten domestic cats. PCR amplifications were carried out with 5 ng of template DNA, 0.3 μM of each primer, 1 x of “11.1 x buffer” [25], 0.06 μl per 10 μl reaction volume of a 20:1 mix of PCRBIO Taq Polymerase (PCR Biosystems; 5 U/μl): Pfu polymerase (2.5 U/μl), with a 10 or 20 μl total volume used for ∼5-kb and ∼8-kb reactions respectively. Primer pairs for the four amplicons are shown schematically in Figure 1b, and sequences and cycling conditions provided in Table S2.

PCR products were visualised by gel electrophoresis. In each case, equimolar concentrations of the respective products for NUMT_1 and 2; NUMT_3 and 4 were pooled together, for each of the ten individuals. Two numt amplicons were sequenced at any one time. DNA libraries were prepared using the native barcoding ligation sequencing genomic DNA protocol for Flongle (version: NBE_9065_v109_revY_14Aug2019) with the NEBNext FFPE DNA repair mix and buffer omitted during the DNA repair and end-prep stage. Each of the two sequencing runs was undertaken using Flongle flow cells with MinKNOW software (v20.10.3).

### Processing of Nanopore sequence data

Guppy (v.5.0.16) was used for basecalling using the high accuracy model with Qscore filtering disabled, and for demultiplexing of Nanopore library native barcodes. Demultiplexing was performed based on the presence of the native barcode on both ends of a read for the MinION flow cell run, and based on the presence of one barcode with Flongle sequencing data (unless stated otherwise). Read quality statistics were assessed using NanoPlot (v.1.33.1) and reads filtered using NanoFilt (v2.8.0) [26].

In mitochondrial sequence analysis, demultiplexed reads were filtered to retain those reads with a quality score of ≥10, and between 100 and 700 bp in length. Primer sequences were removed using Porechop (0.2.4) (github.com/rrwick/Porechop) from the first and last 70 bp of each read, with the `--no_split` function enabled. Trimmed reads from each individual were then mapped to the reference sequence using minimap2 (v2.17) [27]. Output SAM files were transformed to BAM files using SAMtools (v1.13) [28]. Variants were called using freebayes (v0.9.21) [29] with a ploidy of 1. SNVs were retained if they had both a read depth and quality score of ≥ 200, or a quality score of over 100 if the read depth was below this threshold. Consensus sequences were created using BCFtools [30].

Samples sequenced on Flongle in the missing cat case were processed with a few adjustments. Guppy was used for demultiplexing based on the presence of barcodes on both ends of a given read. Reads were filtered using NanoFilt [26] to select reads with a minimum quality of 7 and between 100 and 700 bp in length. Primer sequences were trimmed, and reads were mapped as above. Variant calling was performed using freebayes [29] with the options `-p 1 --min-alternate-fraction 0.5.` Consensus sequences were generated using BCFtools and the resulting sequences were combined with the other 120 domestic cat sequences (119 from ref. [20] together with NC_001700.1). The sequences were aligned using MUSCLE [31].

For numt analysis, reads from both sequencing runs were combined. Following demultiplexing, reads were filtered to remove reads with a quality score <10, and below 3000 or above 17,000 bp in size. Porechop (0.2.4) (github.com/rrwick/Porechop) was used to remove primer sequences from read ends, with the `--no_split` function enabled. The resulting reads were mapped to the reference mitochondrial genome (NC_001700) using minimap2 (v2.17) [27]. SAMtools (v1.13) [28] was used to generate BAM files from the mapping output.

### Data analysis

Sequence data were visualised using IGV, the Integrative Genomics Viewer [32]. Circular representations of mtDNA features and summary data were drawn using shinyCircos [33]. Nomenclature of the basal universal mitotypes A-L and OL-1 was based on variation within the 402-bp CR segment and followed a previously established system [13] with additional UK-specific variants [3]. Mixtures of mitogenome and numt reads were resolved in IGV using the “cluster (phase) alignments” option for each amplicon within the coordinates uniquely spanned by each (see Table S4 for locations).

## Results

### Multiplex mitogenome amplification and sequencing from hair and blood DNA

We designed primers to amplify the whole mitogenome in sixty fragments (average length ∼360 bp) as two sets of thirty non-overlapping amplicons in two separate multiplex PCRs. To test this multiplex, it was applied to matched hair and blood samples from two domestic cats, and products were sequenced on a Nanopore flow cell. Sequencing yielded an average of 869,602 reads per sample, with a minimum read depth of 35 for any base within the mitogenome. We observe a stronger negative correlation between read-depth and amplicon length for hair DNA compared to blood DNA (Figure S1), showing the effect of degradation on the former and supporting the necessity of the short-amplicon multiplex approach.

Figure 2 summarises the short-amplicon multiplex results for matched blood and hair DNA samples from the two cats, and includes long-amplicon data from the same individuals [20] for comparison. There are 144 variants detected for catB29 (Figure 2a), and 151 for catB30 (Figure 2b) using the long-amplicon approach. Variants in both cats are spread unevenly around the mtDNA molecule, with lower representation within the rRNA genes, a finding that most likely represents relatively strong conservation of these sequences, as reflected by partitioned mutation rates in African ape and human data [34], in which rRNA gene mutation rates are similar to rates for the 1st and 2nd codon positions of protein-coding genes. The observed variants match completely with the variants detected in the respective hair DNA, supporting the validity of the multiplex approach for this source. The results for blood DNA, however, are different over the region that corresponds to the major feline numt. Here, an additional 175 variants are observed in catB29 (Figure 2a) which match the published numt DNA variants whilst a further 15 expected mitogenome variants seen in the long-range PCR and hair multiplex were masked by numt bases that also match the mtDNA reference sequence (see Table S3). This indicates that the short-amplicon approach to mitogenome analysis may require deconvolution of the mitogenome and numt components in tissues that contain high levels of nuclear DNA. Analysis in the second domestic cat (catB30; Figure 2b) also reveals numt-specific variants in blood DNA, but far fewer than in catB29. This difference between individuals could reflect different numt copy numbers [12, 19], combined with our majority-nucleotide approach to base calling. Visualisation in IGV with the “cluster (phase) alignments” option shows that for each amplicon separation of the numt and true mtDNA sequences is straightforward due to the high degree of divergence between them (Figure S2), and using this we see that the proportion of numt to mitogenome reads varies between 80-26% (mean 59%) for catB29 and from 47-6% (mean 27%) for catB30 across 23 primer pairs which target mitogenome regions corresponding to the numt and have sufficient homology to permit numt amplification. The proportion of numt-derived reads for each of the 23 amplicons is strongly correlated between the two cats (r_s_= 0.905), with the highest numt penetrance shown by amplicon 22 and the lowest by amplicon 24 in both cats. A further five amplicons in this region (19, 20, 25, 27 & 29) show no evidence of numt bleed-through due to a higher degree of divergence between primer target sites (see Table S4 for variation of numt penetrance by amplicon and individual).

**Figure 2:**
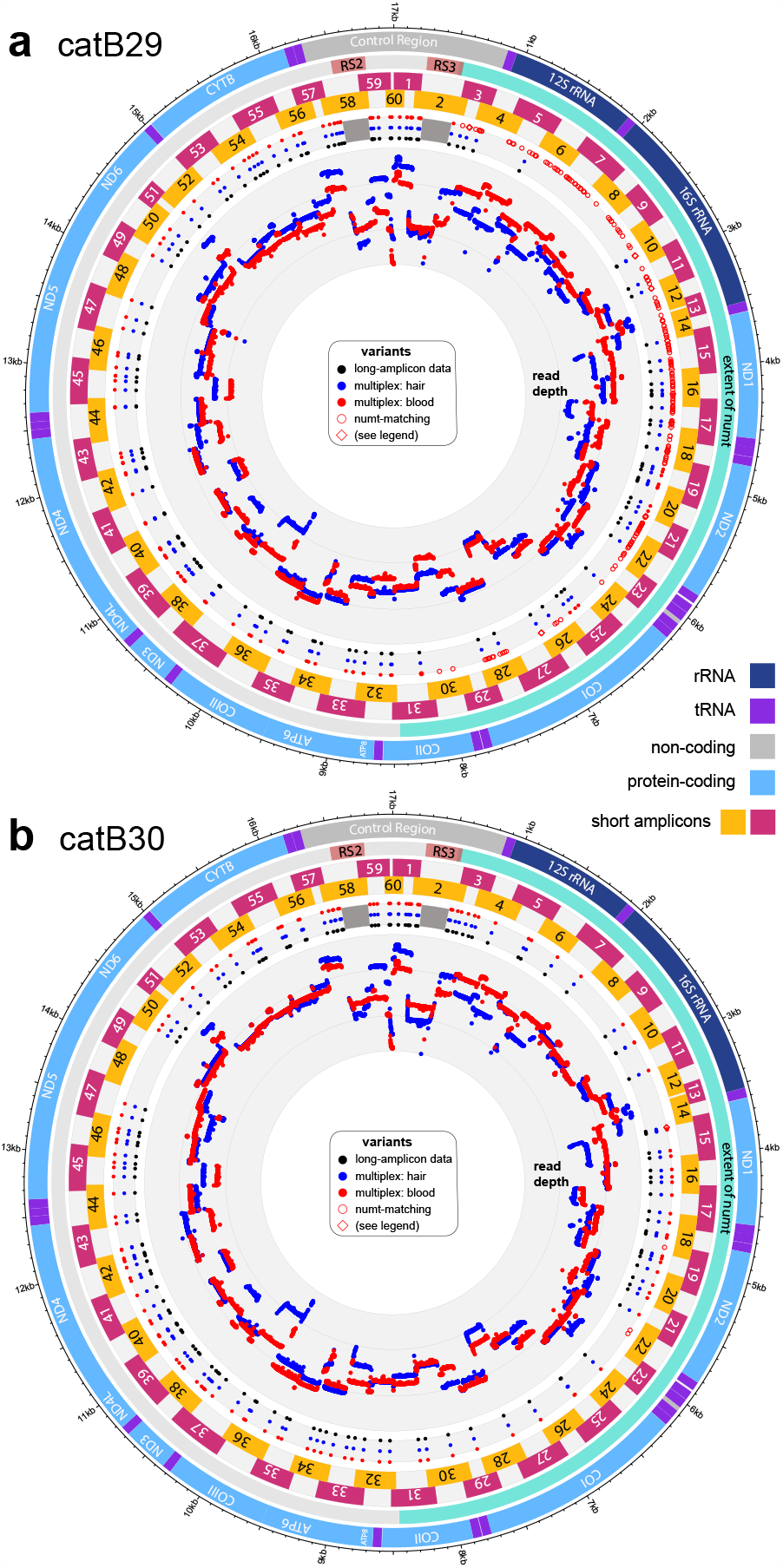
Multiplex sequencing of domestic cat hair avoids numt-specific variants compared to matched blood DNA. Data for two cats are shown: a) catB29, and b) catB30. Locations of the 60 short amplicons are indicated, as well as (inner ring) log_10_ sequence depth for blood (red) and hair DNA (blue). Variants in blood DNA (red filled circles), hair DNA (blue) and in long-amplicon sequence data from blood DNA from the same cat (black) are indicated. Note the complete correspondence between variants detected in long-amplicon data and the hair-based variants, but the excess of blood variants in catB29 and their positional correspondence with the extent of the 7.9-kb numt. Open circles indicate likely numt-specific sequence variants present in the blood DNA data but different from both the mtDNA reference sequence and the analysed cat’s mitogenome. Squares represent numt-specific variants that match the mtDNA reference sequence but differ from the mitogenome sequence of the analysed cat.

### Structures and diversity of numt sequences

Since numt sequences are a potential source of artefacts that could complicate or confound mtDNA analysis [35], we used long-amplicon Nanopore sequencing of numt arrays to better understand numt repeat diversity and its potential to influence the detection of artefacts. We also surveyed recent long-read cat genome sequence data for evidence on the structures of numt arrays.

We designed a long-amplicon approach using numt-specific primers based on the reference sequence (U20754 [19]). These generated four overlapping amplicons ranging from 4.9 to 7.9 kb (Figure 1b) which should allow amplification of adjacent numt copies across the array. This system was used to amplify numt sequences from blood DNAs of ten domestic cats and the products Nanopore-sequenced. The sequenced individuals all matched the NC_001700 mtDNA reference rather than the U20754 numt reference at six SNP sites and four indels (Table S3). It is possible that these reflect numt reference sequence errors, as suggested previously [20] for the cat mtDNA reference sequence (NC_001700; [12]), but they could also represent true variants that are unobserved in our small dataset.

Four publicly available domestic cat (*F. catus*) genome assemblies incorporate data from PacBio long read sequencing. In one assembly [36] the numt array appears to be absent (an earlier study [37] on short-read data from the same cat, Cinnamon, indicated that an array does exist in this genome, but it could not be reliably assembled). The other three assemblies usefully illuminate numt array structures (Figure 3a). Two, derived from hybrid individuals (Fca-508; *F. catus* x *Prionailurus bengalensis* [38] and Fca-126; *F. catus* x *Leopardus geoffroyi*; Bioproject PRJNA773801), contain complete numt arrays consisting of seven to eight repeats of a complete 7.9-kb numt followed by a truncated repeat with an inverted partial repeat at the 3’ end (Figure 3a). The third assembly (AnAms1.0) derived from Senzu, an American shorthair cat [39], includes a partial numt array containing flanking DNA on one side only and 15 near full-length copies of the numt repeat followed by a final truncated inverted copy of the numt (∼3 kb) carrying the same internal deletion as Fca-508 (Figure 3a). The AnAms1.0 numt repeats are more variable in both sequence and length than those from the hybrid individuals, and several of those variants were also detected in our sequencing study.

**Figure 3:**
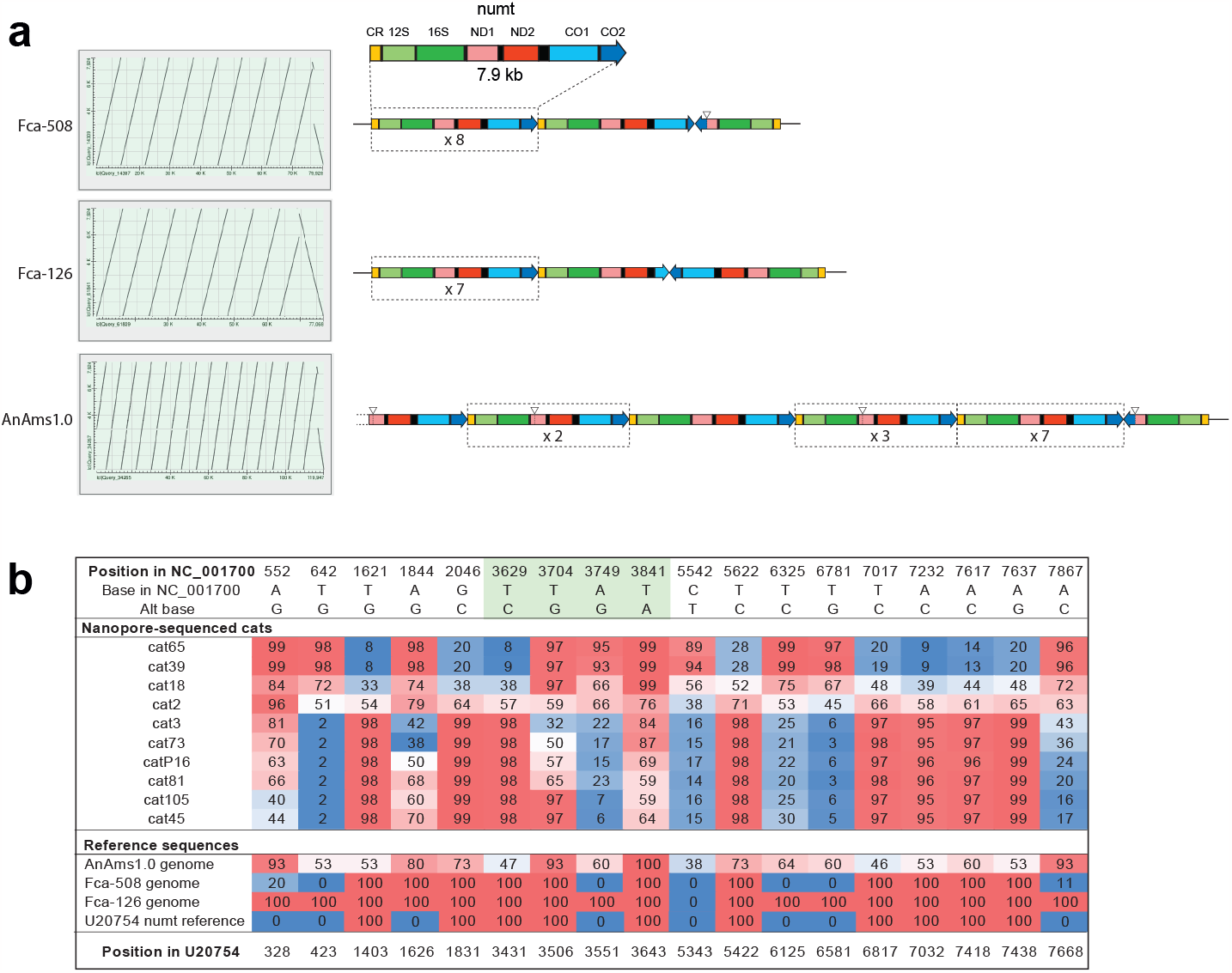
Numt array structures from long-read cat genome assemblies, and variant states in Nanopore-sequenced cats. a) To the left are shown dot-plots generated by aligning the region of the mitogenome (NC_001700:529-8452) corresponding to the span of the numt reference sequence U20754 [19] against three long-read genome assemblies; to the right are the deduced structures of the sequenced arrays. b) Heat-mapped proportions of sequence reads carrying alternative numt bases (derived with respect to the presumed mitogenome sequence) at 18 positions, in ten Nanopore-sequenced cats. Proportions in three reference genomes are also shown. Highlighted in green are four SNVs that illustrate high numt repeat haplotype diversity in phased sequence reads, as shown in Figure S3.

Among the ten analysed cats, 17 sites displayed polymorphic base calls according to the majority variant calling criterion (Figure 3b), with further positions displaying varying proportions of a derived base, although always in the minority of reads (<45%). We found examples where the percentage of variant bases ranged from 0 to >90% among individuals (Figure 3b), presumably affected both by the proportion of variant repeats in the array, the zygosity of the array and the proportion of missing/erroneous base calls due to Nanopore sequencing error. Polymorphic indels were also seen, including the 127-bp deletion observed in half of the AnAms1.0 repeats (corresponding to positions 3430-3557 in the NC_001700 mtDNA reference sequence) which was detected in up to 25% of reads from five cats. Similarly, a 3979-bp deletion spanning positions 3586-7565 was seen in ∼5-20% of reads from all sequenced cats, although any amplification bias toward shorter fragments may cause these deletions to be over-represented. The larger deletion corresponds to that seen in the terminal partial inverted repeat of the AnAms1.0 and Fca-508 arrays. Since all three genome alignments finish with a 3’ terminal inverted repeat, this may be conserved and exist as a single copy in most individuals.

SNP variants tended to be inherited as a haplotype within individual repeats of the numt, but recombination within repeats, as shown by cat81 where all four possible combinations of the two SNP alleles at position 3704 and 3841 are seen, adds to the diversity (Figure 3b; Figure S3). Furthermore, reads were detected that appeared to match mtDNA and could either represent bleed-through of true mitogenome sequence despite the targeting of the primers to numt-specific variants, or evidence that some numt repeats have been converted back to a more mtDNA-like sequence following insertion (Figure S3). As the mtDNA-like sequences were only noted with the one primer pair that could potentially amplify a true mtDNA sequence of <13 kb (i.e. numtF2/R2; Figure 1b) we conclude that these are most likely due to bleed-through of genuine mtDNA. However, some supporting evidence that gene conversion may occur comes from alignment of the AnAms1.0 chr D2 numt repeats against the Lopez mtDNA reference sequence which appears to show patches where conversion has occurred, e.g. in the 358 bases of the terminal AnAms1.0 repeat (AP023162.1:17739103-17739461), there are 24 bases which match the mtDNA reference rather than the numt variants that predominate elsewhere in the array (7151-7544 in NC_001700).

### Matching degraded skeletal remains to hair from a matrilineal relative

Following the development of a multiplex suitable for the analysis of cat hairs (or other equally degraded sample types) typically encountered in forensic scenarios, we had the opportunity to assess its forensic potential in a specific case. The discovery of a cat mandible two months after the disappearance of a female cat led to the owner seeking to establish whether the recovered remains belonged to their missing pet. Mitogenome sequences were obtained from DNA extracted from a mandibular tooth and from a hair sample collected from the missing cat’s male offspring. In order to consider whether a match between the mandible and the offspring would be of value, we considered the probability of a match between the two samples by chance alone. Limiting our comparison to the commonly sequenced 402-bp segment of the control region revealed that both samples belonged to mitotype B, one of the three most common mitotypes (Figure 1d) shared by 23/119 cats in our study. The mitogenome sequences (excluding the RS repetitive regions) of the hair and tooth samples matched perfectly, and were additionally shared with 2/119 cats. The random match probability (RMP) in this case with the CR 402-bp segment was 0.21 and this decreased to 0.03 when considering the whole mitogenome. The decrease in the RMP reflects the limited discrimination power of the widely used short section of the CR and demonstrates the increased power achieved when using the whole mitogenome sequence.

## Discussion

Here we have shown that short-amplicon multiplex amplification and Nanopore sequencing of mitogenomes from cat hairs can provide reliable sequence data that match long-amplicon data from blood taken from the same individuals. Such analysis should have utility in maximising the informativeness of cat hair evidence in forensic cases, by exploiting the ∼9-fold reduction in random match probability that mitogenome sequences provide compared to standard CR analysis [20]. A test case in which degraded DNA from skeletal remains are matched to hair from a matrilineal relative provides proof of principle that the method will be useful in casework. Although we chose to use Nanopore sequencing here, PCR products generated by the multiplex system are also suited to analysis via alternative massively parallel sequencing technologies such as Ion Torrent (Thermo Fisher) or sequencing-by-synthesis (Illumina).

Numts are now known to be widespread in genomes across a wide range of species [40], but were first described in the domestic cat [19]. Applying the mitogenome multiplex to blood DNA reveals additional variants compared to hair DNA (Figure 2), originating from the feline major numt array [19]. This is expected, and analogous results have been observed in extensive studies of human mtDNA data generated with mini-amplicon approaches [35]. However, the extent to which such numt-derived variants manifest varies greatly between the two cats analysed here, and indeed between different amplicons (Figure 2), which may reflect differences in numt copy number and sequence. To understand the possible influence of numt sequences in tissue types where mixed nuclear and mitochondrial sequences might occur, we mined three available long-read cat genome assemblies ([38, 39]; Bioproject PRJNA773801) and also analysed sequence variation in numt arrays via long-amplicon Nanopore sequencing.

The original report on the feline numt [19] estimated array size by pulsed-field gel electrophoresis of BglII-restriction-digested cat DNAs, followed by Southern blotting and hybridisation with a numt probe. In three pedigrees encompassing a total of nine independent numt alleles, this gave fragment sizes of 300-600 kb, interpreted as equivalent to 38-76 copies of a 7.9 kb sequence. However, the two independent and complete numt arrays now available from long-read sequencing (Figure 3a) each contain only nine repeats, well below the 38-repeat lower limit suggested previously [19]. Bioinformatic location of the closest BglII sites flanking the Fca-126 and Fca-508 arrays shows that the numt-containing BglII fragment in each case contains 50 kb of flanking DNA in addition to the numt array itself. Accounting for this non-numt material reduces the published numt copy number somewhat, from 38-76 to 31-69 repeats; however, this range still greatly exceeds the values observed here in sequenced genomes. Given that the AnAms1.0 numt array is incomplete, that example may well lie in the range originally described [19].

Further work is needed, either via long-read assemblies or qPCR, to more accurately assess numt copy number and sequence variation in cat populations. As a first step, we approached the issue in a simple way by designing an overlapping numt-specific long-amplicon system (Figure 1b), followed by the bioinformatic sorting of reads with the help of phased variants that are linked within a given read ([35]; Figure S3). Using this approach, we identified a proportion of reads in all cats that encompassed the 3979-bp deletion seen only in the single inverted terminal repeat of the numt array in two of the genome assemblies. If all numt arrays terminated in a single copy of this inverted repeat, then we could estimate the number of other repeats from the proportion of reads carrying this deletion. Based on this, the numt Nanopore sequence data from ten cats would suggest that the copy number of near complete repeats at the 5’ end of the array ranges from about 5 to 20, encompassing the observed numbers in the long-read sequenced arrays (Figure 3a); however, it is likely that PCR amplification bias towards any shorter repeat units causes underestimation of the true number of full length repeats. As evidenced by the Fca-126 assembly, a minority of numt arrays in our dataset may lack the 3979-bp deletion in the inverted terminal repeat and therefore would not be distinguishable from the normally orientated repeats using our PCR assay: this would lead to overestimation of the total repeat number in cats with this terminal repeat.

In multiplex analysis of blood DNA, the proportion of reads originating from the numt rather than the mitogenome is affected not only by numt copy number but also by the ability of the multiplex primers to amplify the diverged numt sequence. We did not observe numt-specific variants when multiplex primer pairs included between one to five mismatches with the numt sequence, and where at least one of these mismatches was present within three bases of the 3’ end. As a consequence, the penetrance of the numt array also varies from amplicon to amplicon. Despite the many sources of uncertainty in the proportion of reads that may have a numt origin they can easily be bioinformatically identified and excluded from mitogenome analysis using phasing in IGV and the numt sequence data presented here. Through these means it is now possible to discriminate between individual cats to a much greater degree when considering cat hairs or other degraded DNA sources as evidential material.

## Data availability

Genbank accession numbers for mitogenome sequences are: OR077846 (catB29) and OR095104 (catB30). Sequence information for numt amplicons can be found in the Sequence Read Archive under BioProject PRJNA981977.

## Acknowledgments

EP was supported by a BBSRC-MIBTP (grant no. BB/M01116X/1) iCASE studentship, co-sponsored by Twycross Zoo (East Midland Zoological Society) and Zoological Society of London. We thank Lisa Gillespie for assistance, and Jane and Andrew Elliot for additional financial support. This research used the SPECTRE High Performance Computing Facility at the University of Leicester for data analysis.

## References

[1] Cats Protection, Cats Report, https://www.cats.org.uk/about-cp/cats-report-2022, 2022.

[2] L.A. Lyons, R.A. Grahn, T.J. Kun, L.R. Netzel, E.E. Wictum, J.L. Halverson, Acceptance of domestic cat mitochondrial DNA in a criminal proceeding, Forensic Sci Int Genet 13 (2014) 61–7.

[3] B. Ottolini, G.M. Lall, F. Sacchini, M.A. Jobling, J.H. Wetton, Application of a mitochondrial DNA control region frequency database for UK domestic cats, Forensic Sci Int Genet 27 (2017) 149–155.

[4] R.A. Grahn, H. Alhaddad, P.C. Alves, E. Randi, N.E. Waly, L.A. Lyons, Feline mitochondrial DNA sampling for forensic analysis: when enough is enough!, Forensic Sci Int Genet 16 (2015) 52–57.

[5] C.R. Tarditi, R.A. Grahn, J.J. Evans, J.D. Kurushima, L.A. Lyons, Mitochondrial DNA sequencing of cat hair: an informative forensic tool, Journal of forensic sciences 56 Suppl 1 (2011) S36–46.

[6] M.A. Menotti-Raymond, V.A. David, L.L. Wachter, J.M. Butler, S.J. O’Brien, An STR forensic typing system for genetic individualization of domestic cat (Felis catus) samples, J Forensic Sci 50 (2005) 1061–70.

[7] M.J. Lipinski, Y. Amigues, M. Blasi, T.E. Broad, C. Cherbonnel, G.J. Cho, S. Corley, P. Daftari, D.R. Delattre, S. Dileanis, J.M. Flynn, D. Grattapaglia, A. Guthrie, C. Harper, P.L. Karttunen, H. Kimura, G.M. Lewis, M. Longeri, J.C. Meriaux, M. Morita, C. Morrin-O’Donnell R, T. Niini, N.C. Pedersen, G. Perrotta, M. Polli, S. Rittler, R. Schubbert, M.G. Strillacci, H. Van Haeringen, W. Van Haeringen, L.A. Lyons, An international parentage and identification panel for the domestic cat (Felis catus), Animal genetics 38 (2007) 371–7.

[8] A. Linacre, Animal Forensic Genetics, Genes (Basel) 12 (2021).

[9] A. Brooks, E.K. Creighton, B. Gandolfi, R. Khan, R.A. Grahn, L.A. Lyons, SNP Miniplexes for Individual Identification of Random-Bred Domestic Cats, Journal of forensic sciences 61 (2016) 594–606.

[10] C.A. Linch, Degeneration of nuclei and mitochondria in human hairs, Journal of forensic sciences 54 (2009) 346–9.

[11] B. Budowle, M.W. Allard, M.R. Wilson, R. Chakraborty, Forensics and mitochondrial DNA: applications, debates and foundations, Ann Rev Genomics Hum Genet 4 (2003) 119–141.

[12] J.V. Lopez, S. Cevario, S.J. O’Brien, Complete nucleotide sequences of the domestic cat (Felis catus) mitochondrial genome and a transposed mtDNA tandem repeat (Numt) in the nuclear genome, Genomics 33 (1996) 229–46.

[13] R.A. Grahn, J.D. Kurushima, N.C. Billings, J.C. Grahn, J.L. Halverson, E. Hammer, C.K. Ho, T.J. Kun, J.K. Levy, M.J. Lipinski, J.M. Mwenda, H. Ozpinar, R.K. Schuster, S.J. Shoorijeh, C.R. Tarditi, N.E. Waly, E.J. Wictum, L.A. Lyons, Feline non-repetitive mitochondrial DNA control region database for forensic evidence, Forensic Sci Int Genet 5 (2011) 33–42.

[14] I. Glazewska, T. Kijewski, A new view on the European feline population from mtDNA analysis in Polish domestic cats, Forensic Sci Int Genet 27 (2017) 116–122.

[15] M. Arcieri, G. Agostinelli, Z. Gray, A. Spadaro, L.A. Lyons, K.M. Webb, Establishing a database of Canadian feline mitotypes for forensic use, Forensic Sci Int Genet 22 (2016) 169–74.

[16] M. Wesselink, S. Desmyter, I. Kuiper, Local populations and inaccuracies: Determining the relevant mitochondrial haplotype distributions for North West European cats, Forensic Sci Int Genet 30 (2017) 71–80.

[17] M. Wesselink, L. Bergwerff, D. Hoogmoed, A.D. Kloosterman, I. Kuiper, Forensic utility of the feline mitochondrial control region - A Dutch perspective, Forensic Sci Int Genet 17 (2015) 25–32.

[18] J.L. Halverson, C. Basten, Forensic DNA identification of animal-derived trace evidence: tools for linking victims and suspects, Croatian medical journal 46 (2005) 598–605.

[19] J.V. Lopez, N. Yuhki, R. Masuda, W. Modi, S.J. O’Brien, Numt, a recent transfer and tandem amplification of mitochondrial DNA to the nuclear genome of the domestic cat, J Mol Evol 39 (1994) 174–90.

[20] E. Patterson, G. Matharu Lall, R. Neumann, B. Ottolini, C. Batini, F. Sacchini, A.P. Foster, J.H. Wetton, M.A. Jobling, Mitogenome sequences of domestic cats demonstrate lineage expansions and dynamic mutation processes in a mitochondrial minisatellite, BioRxiv 544779 (2023) https://doi.org/10.1101/2023.06.13.544779.

[21] L. Hasap, W. Chotigeat, J. Pradutkanchana, U. Vongvatcharanon, T. Kitpipit, P. Thanakiatkrai, A novel, 4-h DNA extraction method for STR typing of casework bone samples, Int J Legal Med 134 (2020) 461–471.

[22] A. Larsson, AliView: a fast and lightweight alignment viewer and editor for large datasets, Bioinformatics 30 (2014) 3276–8.

[23] P.M. Vallone, J.M. Butler, AutoDimer: a screening tool for primer-dimer and hairpin structures, Biotechniques 37 (2004) 226–31.

[24] J. Ye, G. Coulouris, I. Zaretskaya, I. Cutcutache, S. Rozen, T.L. Madden, Primer-BLAST: a tool to design target-specific primers for polymerase chain reaction, BMC bioinformatics 13 (2012) 134.

[25] A.J. Jeffreys, R. Neumann, V. Wilson, Repeat unit sequence variation in minisatellites: a novel source of DNA polymorphism for studying variation and mutation by single molecule analysis, Cell 60 (1990) 473–485.

[26] W. De Coster, S. D’Hert, D.T. Schultz, M. Cruts, C. Van Broeckhoven, NanoPack: visualizing and processing long-read sequencing data, Bioinformatics 34 (2018) 2666–2669.

[27] H. Li, Minimap2: pairwise alignment for nucleotide sequences, Bioinformatics 34 (2018) 3094–3100.

[28] H. Li, B. Handsaker, A. Wysoker, T. Fennell, J. Ruan, N. Homer, G. Marth, G. Abecasis, R. Durbin, The Sequence Alignment/Map format and SAMtools, Bioinformatics 25 (2009) 2078–2079.

[29] E. Garrison, G. Marth, Haplotype-based variant detection from short-read sequencing, arXiv 1207.3907 (2012).

[30] P. Danecek, A. Auton, G. Abecasis, C.A. Albers, E. Banks, M.A. DePristo, R.E. Handsaker, G. Lunter, G.T. Marth, S.T. Sherry, G. McVean, R. Durbin, The variant call format and VCFtools, Bioinformatics 27 (2011) 2156–2158.

[31] R.C. Edgar, MUSCLE: multiple sequence alignment with high accuracy and high throughput, Nucleic Acids Res 32 (2004) 1792–7.

[32] J.T. Robinson, H. Thorvaldsdottir, W. Winckler, M. Guttman, E.S. Lander, G. Getz, J.P. Mesirov, Integrative genomics viewer, Nat Biotechnol 29 (2011) 24–26.

[33] Y. Yu, Y. Ouyang, W. Yao, shinyCircos: an R/Shiny application for interactive creation of Circos plot, Bioinformatics 34 (2018) 1229–1231.

[34] P. Soares, L. Ermini, N. Thomson, M. Mormina, T. Rito, A. Rohl, A. Salas, S. Oppenheimer, V. Macaulay, M.B. Richards, Correcting for purifying selection: an improved human mitochondrial molecular clock, Am J Hum Genet 84 (2009) 740–759.

[35] C. Marshall, W. Parson, Interpreting NUMTs in forensic genetics: Seeing the forest for the trees, Forensic Sci Int Genet 53 (2021) 102497.

[36] R.M. Buckley, B.W. Davis, W.A. Brashear, F.H.G. Farias, K. Kuroki, T. Graves, L.W. Hillier, M. Kremitzki, G. Li, R.P. Middleton, P. Minx, C. Tomlinson, L.A. Lyons, W.J. Murphy, W.C. Warren, A new domestic cat genome assembly based on long sequence reads empowers feline genomic medicine and identifies a novel gene for dwarfism, PLoS Genet 16 (2020) e1008926.

[37] A. Antunes, J. Pontius, M.J. Ramos, S.J. O’Brien, W.E. Johnson, Mitochondrial introgressions into the nuclear genome of the domestic cat, J Hered 98 (2007) 414–20.

[38] K.R. Bredemeyer, A.J. Harris, G. Li, L. Zhao, N.M. Foley, M. Roelke-Parker, S.J. O’Brien, L.A. Lyons, W.C. Warren, W.J. Murphy, Ultracontinuous Single Haplotype Genome Assemblies for the Domestic Cat (Felis catus) and Asian Leopard Cat (Prionailurus bengalensis), J Hered 112 (2021) 165–173.

[39] S. Isobe, Y. Matsumoto, C. Chung, M. Sakamoto, T.F. Chan, H. Hirakawa, G. Ishihara, H.M. Lam, S. Nakayama, S. Sasamoto, Y. Tanizawa, A. Watanabe, K. Watanabe, M. Yagura, Y. Nakamura, AnAms1.0: A high-quality chromosome-scale assembly of a domestic cat Felis catus of American Shorthair breed, BioRxiv (2020) doi.org/10.1101/2020.05.19.103788.

[40] F.M. Calabrese, D.L. Balacco, R. Preste, M.A. Diroma, R. Forino, M. Ventura, M. Attimonelli, NumtS colonization in mammalian genomes, Sci Rep 7 (2017) 16357.

[41] A.R. Hoelzel, J.V. Lopez, G.A. Dover, S.J. O’Brien, Rapid evolution of a heteroplasmic repetitive sequence in the mitochondrial DNA control region of carnivores, J Mol Evol 39 (1994) 191–9.

[42] J.H. Kim, A. Antunes, S.J. Luo, J. Menninger, W.G. Nash, S.J. O’Brien, W.E. Johnson, Evolutionary analysis of a large mtDNA translocation (numt) into the nuclear genome of the Panthera genus species, Gene 366 (2006) 292–302.

