## Supplementary material for "Defining cat mitogenome variation and accounting for numts via multiplex amplification and Nanopore sequencing": Figures S1-S3

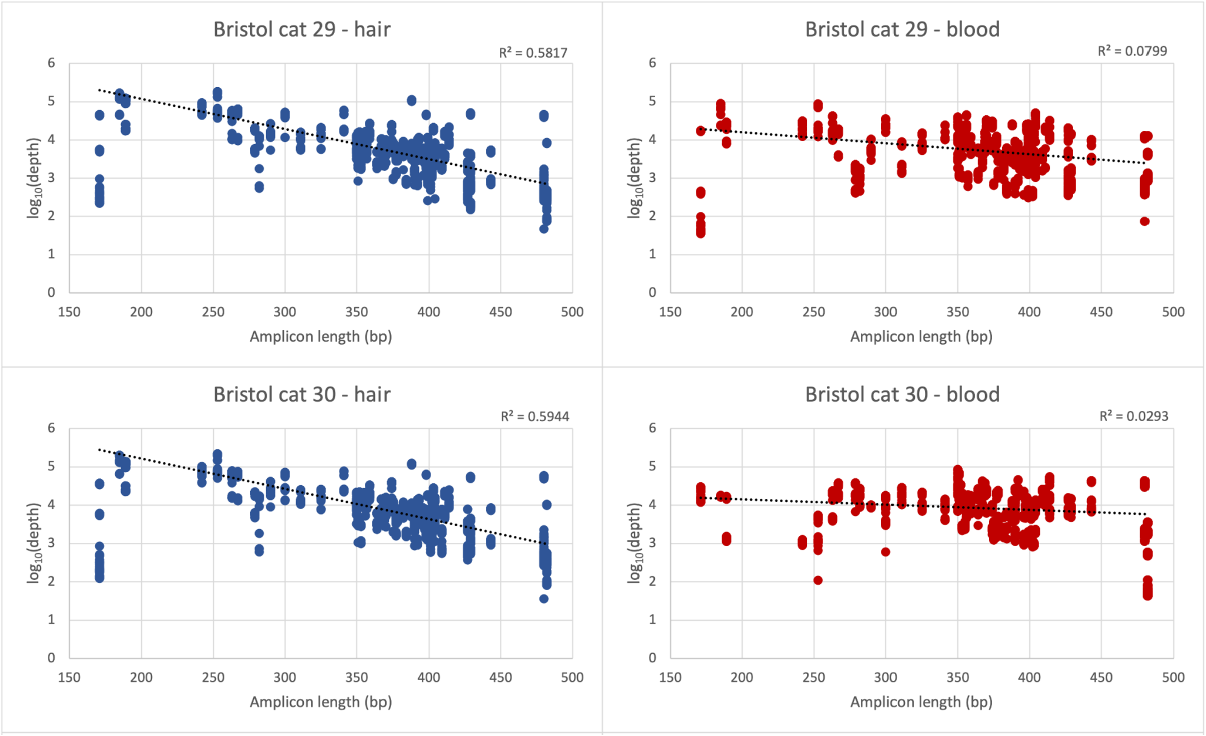


**Figure S1: Relationship between $\log_{10}$ sequence read depth and amplicon length for matched hair and blood DNA samples in two domestic cats.**

Read depth declines more steeply with increasing amplicon length in hair than in blood, reflecting the relatively degraded nature of hair mtDNA.

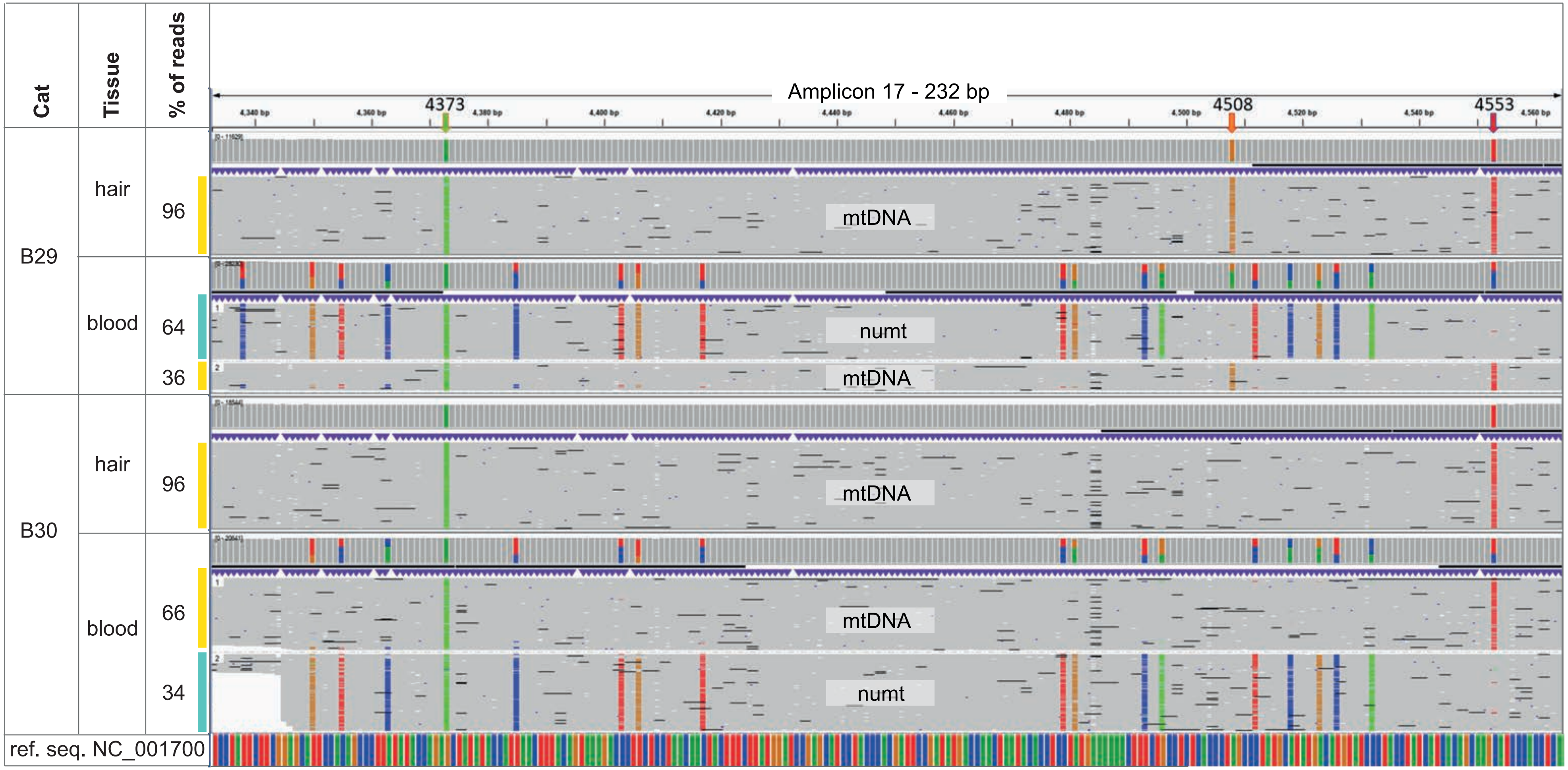


**Figure S2: IGV screenshot of matched hair and blood multiplex amplifications.**

An example of IGV clustering of phased variants for amplicon 17 encompassing three mitogenome SNV sites - 4373G/A, 4508A/G & 4553G/T in multiplex amplifications of cats B29 and B30. The 4373A allele is fixed in all 119 typed mitogenomes and sequenced numt arrays suggestive of an error in the NC_001700 reference sequence, 4508G is a defining variant for haplogroup A (which includes catB29), and 4553T is the derived variant shared by all haplogroups except H (to which the cat reference NC_001700 belongs). Phasing detects only a single haplotype in hair comprising >95% of reads, whereas catB29 blood shows a majority of numt variant calls (64%) across 18 numt defining sites and catB30 blood a majority of mitogenome reads (66%).

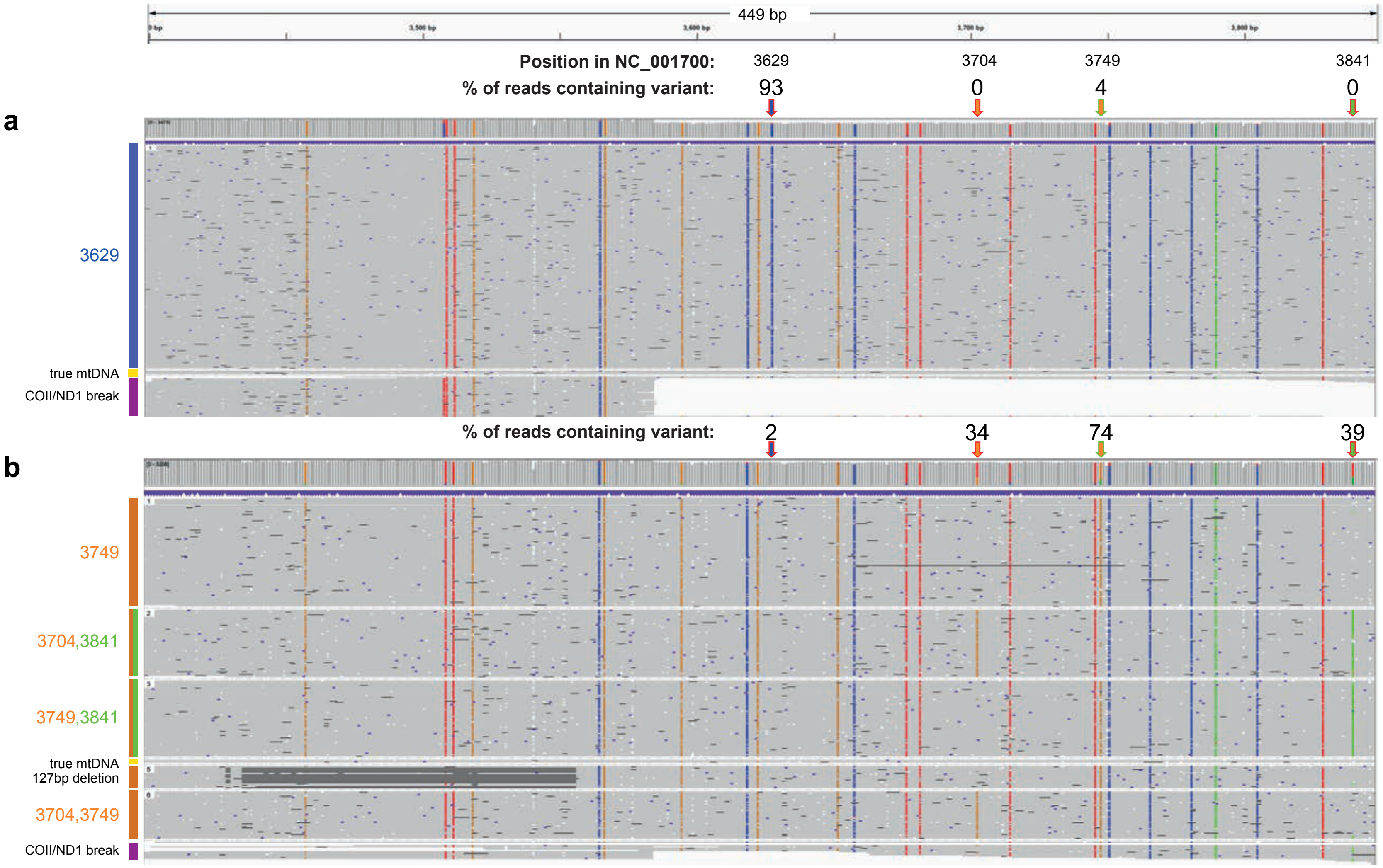


**Figure S3: IGV screenshots showing examples of homogeneous and heterogeneous numt arrays in two domestic cats.**

IGV screenshots illustrating clusters of phased variants in the most diverse region of numt repeats (covering the four SNV sites highlighted green in Figure 3b and two frequent large deletions) a) a low-diversity numt (cat65) displaying a single uninterrupted repeat type, a high proportion of reads terminating at the breakpoint that characterises the inverted terminal repeat (possibly indicating a short total array length) and a small number of bleed-through mtDNA reads which match the NC_001700 mtDNA reference sequence and hence are devoid of variants (vertical coloured bars). b) a high-diversity numt (cat81) with several distinct repeat types including repeats encompassing the 127-bp deletion and a lower proportion of reads originating from the inverted terminal repeat.
